# AIRR-C Human IG Reference Sets: curated sets of immunoglobulin heavy and light chain germline genes

**DOI:** 10.1101/2023.09.01.555348

**Authors:** Andrew M. Collins, Mats Ohlin, Martin Corcoran, James M. Heather, Duncan Ralph, Mansun Law, Jesus Martínez-Barnetche, Jian Ye, Eve Richardson, William S. Gibson, Oscar L. Rodriguez, Ayelet Peres, Gur Yaari, Corey T. Watson, William D. Lees

**Author notes:** Corresponding authors: Andrew M. Collins William D. Lees.

## Abstract

Analysis of an individual’s immunoglobulin (IG) gene repertoire requires the use of high-quality germline gene Reference Sets. The Adaptive Immune Receptor Repertoire-Community (AIRR-C) Reference Sets have been developed to include only human IG heavy and light chain alleles that have been confirmed by evidence from multiple high-quality sources. By including only those alleles with a high level of support, including some new sequences that currently lack official names, AIRR-seq analysis will have greater accuracy and studies of the evolution of immunoglobulin genes, their allelic variants and the expressed immune repertoire will be facilitated. Although containing less than half the previously recognised IG alleles (e.g. just 198 IGHV sequences), the Reference Sets eliminated erroneous calls and provided excellent coverage when tested on a set of repertoires from 99 individuals comprising over 4 million V(D)J rearrangements. To improve AIRR-seq analysis, some alleles have been extended to deal with short 3’ or 5’ truncations that can lead them to be overlooked by alignment utilities. To avoid other challenges for analysis programs, exact paralogs (e.g. IGHV1-69*01 and IGHV1-69D*01) are only represented once in each set, though alternative sequence names are noted in accompanying metadata. The Reference Sets also include novel alleles: 8 IGHV alleles, 2 IGKV alleles and 5 IGLV alleles. The version-tracked AIRR-C Reference Sets are freely available at the OGRDB website (https://ogrdb.airr-community.org/germline_sets/Human) and will be regularly updated to include newly-observed and previously-reported sequences that can be confirmed by new high-quality data.

## Introduction

Cellular and humoral immune responses are key components of our defence against external threats from a range of pathogens [1; 2], as well as internal threats like oncogenic transformation and cancer [3]. Repertoire analysis of the B and T cell transcriptome is now a part of many studies of the immune response, and such analyses have deepened our understanding of immune protection against, for example, Human Immunodeficiency Virus-1 [4], influenza viruses [5], and SARS-CoV-2 [6]. Such analyses can guide the development of novel vaccine strategies [7], and can lead to the identification of highly functional antibodies with the potential to be translated into anti-viral drugs of clinical utility [8; 9; 10].

An important aspect of immunoglobulin (IG) repertoire analysis is the identification of germline V, D and J genes that have contributed to each V(D)J gene sequence. Germline gene Reference Sets - made up of known IG allelic variants - are critical for these kinds of analyses. The identification of germline genes that contribute to the formation of V(D)J gene sequences allows the clonal relationships between different sequences to be identified [11; 12; 13; 14]. Analysis of V(D)J rearrangements also allows somatic point mutations within the gene rearrangements to be determined, giving insights into the development of antibody specificities and affinity-driven selection [15; 16; 17] as well as antibody isotype functions [18; 19; 20]. If these kinds of analyses are to be improved, the available Reference Sets must also be improved.

Over the last decade, the number of reported germline sequences has grown substantially as a result of new gene discoveries [21; 22; 23; 24; 25]. It is likely that most common allelic variants that are found in well-studied populations have now been identified, but a large number of variants from less-studied populations probably remain to be found [26; 27; 28].

Rare alleles remain to be documented in all populations [22], and population coverage must be increased [29]. Fortunately, next-generation sequencing of the IG loci should soon provide this population coverage, but challenges associated with new sequencing technologies mean that IG gene discovery must be approached with great care [30; 31].

In recent years most reports of newly-discovered alleles have come from the analysis of AIRR-seq data. Since 2018, 34 inferred IGHV allelic variants, as well as 6 IGLV and 3 IGKV variants, have been validated and assigned temporary names (e.g. IGHV1-69*i04) by the Inferred Allele Review Committee (IARC) of the Adaptive Immune Receptor Repertoire-Community (AIRR-C) [32]. The IARC is now one of two review committees operating at the direction of the International Union of Immunological Societies (IUIS) Immunoglobulin (IG), T cell receptor (TR) and Major Histocompatibility (MH) Nomenclature Sub-committee. The second review committee, the TR-IG Nomenclature Review Committee (TR-NRC), is responsible for the official naming of IG genes. To date, 33 of the 43 inferred sequences affirmed by the IARC have been officially named by IUIS after referral from the IARC [33].

The nomenclature used is referred to here as the IUIS nomenclature to reflect IUIS’s responsibility for naming. It was developed by Marie-Paule Lefranc, the founder of the ImMunoGeneTics (IMGT) group, and the former chair of the IUIS IG, TR and MH Nomenclature Subcommittee [34; 35; 36]. Other sequences that have not yet been assigned official names are referred to here by their temporary IARC names, or by unofficial names indicating their differences from other named sequences (e.g. IGLV3-25*03_G74C).

Although the inferred IG sequences have not been mapped to any reported human genome assembly, their temporary names are modelled on the official IUIS human IG nomenclature – a positional nomenclature. The general principle of the IARC naming process has been that when a putative variant sequence aligns to a known allele of a single gene with high sequence identity, and with substantially lower identity to the alleles of other genes, the sequence can reasonably be assigned as a variant of that gene. In many instances, however, an assignment cannot be made with high certainty based on sequence identity. There may be several similar genes that could all be assigned as the germline gene of the novel allele. The IARC continues to assign temporary names, but their most recent reports make clear that the genomic locations of some new alleles must be considered uncertain.

Despite the growth of historical Reference Sets in recent years [37; 38], many nucleotide and structural level polymorphisms (large insertions, duplications and deletions) remain to be identified in the human population. The ideal Reference Set for the study of an individual’s immune response is therefore a personalised one, determined by genomic sequencing prior to V(D)J sequence annotation. Alternatively, a personalised repertoire developed from AIRR-seq data (expressed V(D)J gene rearrangements) using inference technology can be used for high-quality sequence annotation [11; 39]. In practice, most repertoire studies rely on standard publicly available Reference Sets for their analyses [31; 37; 38], and even the inference of a personalised repertoire begins with analysis using a standard set.

Previously reported reference sets have been compiled from sequences that have been reported over several decades. Over this time, as sequencing technology has changed, the risks of sequencing errors have also changed. Today errors may arise, for example, from the annotation of poor genome assemblies [30]. In the past, sequencing itself was so error-prone that it is likely that many sequences reported in the 1980s and 1990s included sequencing errors [40].

Sufficient published research now exists to create human Reference Sets in which each sequence is validated by at least two high-quality sources, providing confidence that erroneous sequences have been excluded. In this report we describe how well documented alleles that were reported in the past have been combined with newly discovered IGH and IG light chain (lambda, IGL; kappa, IGK) alleles to produce the high-quality AIRR-C IGH_VDJ, IGKappa_VJ and IGLambda_VJ Reference Sets.

The AIRR-C Reference Sets include a small number of sequences that have artificial extensions, for some reported germline sequences appear to have short truncations, usually at their 3’ ends. Extensions of these sequences in the AIRR-C Reference Sets will help ensure that they are not overlooked by sequence alignment utilities, and the presence of extensions is noted in their sequence metadata. The AIRR-C Reference Sets also include modifications to deal with issues arising with sequences that appear in the genome as exact paralogs (duplicates). In AIRR-seq analysis, the presence of identical sequences in the Reference Set can cause downstream analytical pipelines to overlook both copies of the sequence. A single copy of each sequence pair is therefore retained in the Reference Set with the missing sequence being recorded as a paralog in the sequence metadata. A script is available at the OGRDB website that facilitates the production of Reference Sets that have the details of exact paralogs in the FASTA header (https://airr-community.github.io/receptor-germline-tools/_build/html/introduction.html).

The AIRR-C IG Reference Sets are published under the FAIR guiding principles for scientific data management [41], using a recently described schema [33], with a minimally restricted licence. The Reference Sets will grow over time by the inclusion of newly identified allelic variants, as well as by the inclusion of previously reported sequences whose existence may come to be confirmed from the accumulation of new evidence. The nature of these changes to the Reference Sets and evidence in support of those changes will be clearly documented at the OGRDB website (https://ogrdb.airr-community.org) [42]. The AIRR-C Reference Sets will be strictly versioned, and versioned sets will be referenceable via Digital Object Identifiers (DOIs). This will mean that proper documentation in the literature of the use of the Reference Sets will allow readers to easily understand the germline gene data that has been used in an analysis.

## Method

### Evaluation of the effects of short 3’ truncations of Reference sequences upon AIRR-seq analysis

The way that AIRR-seq analysis is influenced by changes in the lengths of sequences in Reference Sets was investigated using transcriptome data sets of project PRJEB26509 from the European Nucleotide Archive [43], generated from FACS-sorted (CD19^+^ CD27^-^ IgD^+^ IgA^-^ IgG^-^) naïve B cells. These datasets can be expected to express high frequencies of unmutated immunoglobulin-encoding V(D)J genes. Six samples were selected for analysis based upon their carriage of different alleles of the IGHV4-38-2 gene. (A different analysis elsewhere in this study involves the complete PRJEB26509 dataset, where it is referred to as the Gidoni-VDJbase dataset.) Reads were assembled using PEAR v0.9.6. Filtering was performed using the default IgDiscover settings that removed reads with no J gene assigned, with stop codons, with V gene coverage <90%, with J gene coverage <60%, and/or with V gene E-value >1E-3. Annotation was performed within the IgDiscover 0.11 framework using IgBLAST, and Reference Sets including an archived IMGT IGHV Reference Set (accessed January 2020) and a modified set that incorporated an extension of IGHV4-38-2*01 by two nucleotides to resolve a truncation of its 3’-end.

### Compilation and evaluation of data relating to IGHV, IGKV and IGLV genes

The AIRR-C IG Reference Sets were produced using a protocol outlined in Figure 1. Sets of candidate V, D and J sequences were first compiled either directly from a number of key reports, or by investigation of GenBank submissions associated with those reports. This included the GRCh38 genome reference sequence and other assemblies produced using similar sequencing and assembly methods. Orphons, which are non-functional IG-like genes found outside the IG loci, were not considered for inclusion. V sequences with apparent 5’ or 3’ truncations of 25 or more nucleotides were also excluded. Candidate sequences were then restricted to those genes for which at least one allele has been reported that is an Open Reading Frame (ORF) and that includes both the first and the second conserved cysteines of the sequence. In this way, many pseudogenes were excluded from consideration. Of the remaining sequences, a candidate V sequence was ultimately included in what is referred to here as a Source Set if it could be confirmed from at least two of these independent reports, or if the reported sequence could be confirmed by additional evidence from particular AIRR-seq datasets that had been evaluated for the quality of the methods used in their generation. The Source Sets were finally used to produce IGH_VDJ, IGKappa_VJ and IGLambda_VJ Reference Sets by the extension of (likely) truncated sequences, and after dealing with exact paralogs as outlined below.

**Figure 1:**
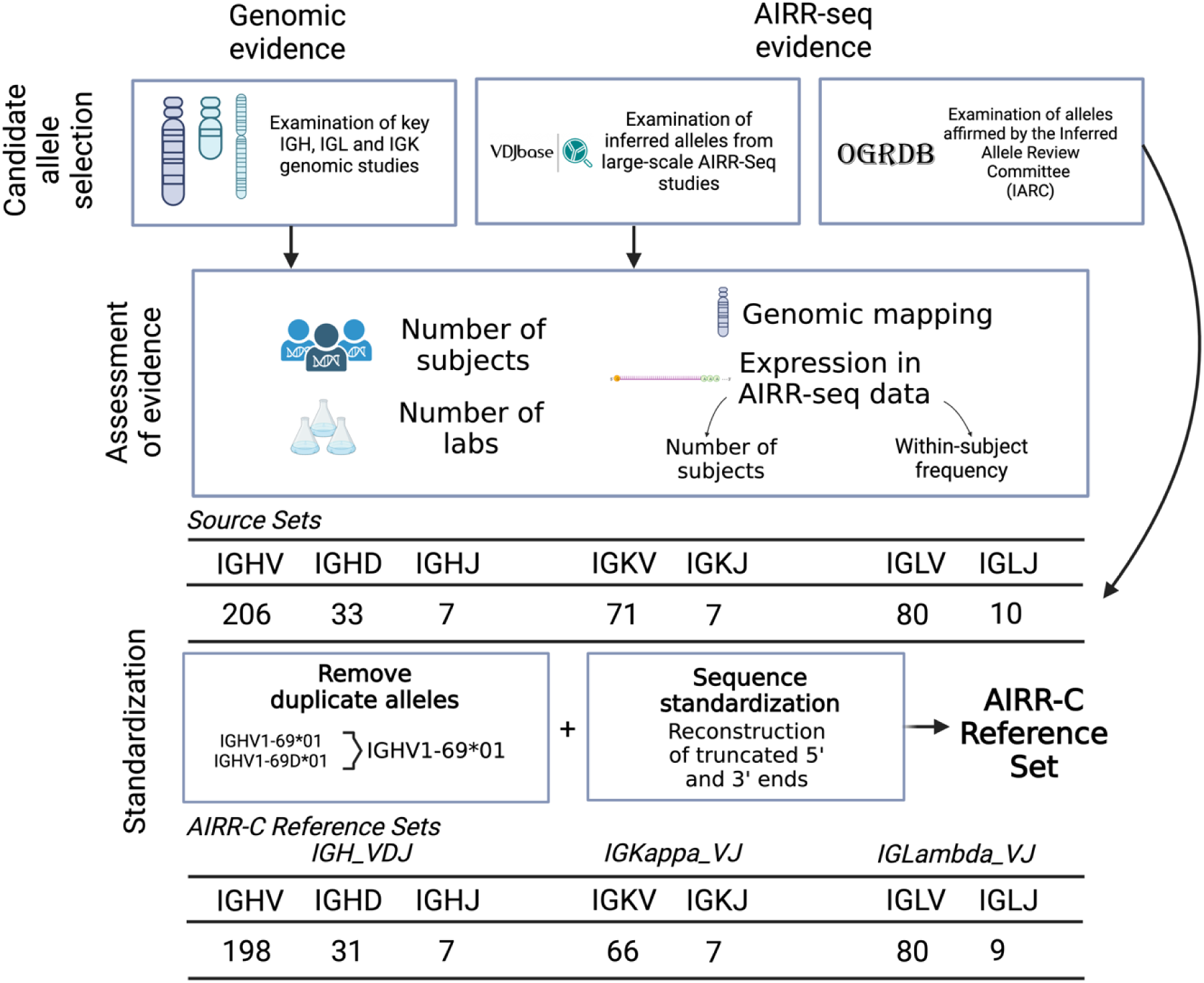
The compilation of Reference Sets began with the identification of candidate sequences from a limited number of key genomic and AIRR-seq studies, as well as reports from the Inferred Allele Review Committee (IARC). Evaluation of evidence in support of each candidate sequence led to the preparation of Source Sets. These include all supported sequences as officially recognized. Reference Sets were then prepared from the Source Sets by retention of just a single copy of duplicated sequences, and by the artificial extension of sequences that appear to be truncated at their 3’ or 5’ ends.

The key reports used to compile the set of candidate IGHV sequences included the historically important genomic studies of the IGH locus by Matsuda and colleagues [44; 45] and Watson and colleagues [25]. The recent genomic study of 154 individuals of diverse ethnicity, by Rodriguez and colleagues, was also included [46]. Candidate IGKV and IGLV sequences were compiled by reference to critical studies from the Zachau group [47; 48; 49], the Winter/Tomlinson group [50; 51; 52], the Watson group [24; 26], and the Shimizu group [53; 54]. Sequences from the key studies were identified directly from the literature or from BLAST searches of GenBank. Genomic IGLV and IGLJ sequences from a recent study [26] were accessed at the VDJbase website (VDJbase.org). In this study, the authors generated haplotyped assemblies of the IGL locus from 16 individuals using a combination of long-read whole-genome sequencing, fosmid technology and capture-probe long-read sequencing. The assemblies were annotated using IGenotyper [55]. The authors also deposited at VDJbase annotations of 32 long-read assemblies created by the Human Pangenome Project [56]. This data is collectively referred to here as the Gibson-VDJbase dataset.

Heavy and light chain sequences identified in the genomic studies of Mikocziova and colleagues [21; 22], genomic studies of IGH genes in African and Melanesian individuals [27; 28] and heavy chain sequences identified by genomic validation (individual D19) in the study of Narang and colleagues [57] were then included as candidate sequences.

Additional IG sequences inferred from AIRR-seq data and affirmed by the IARC were also added to the set of candidate sequences [32]. These IARC-affirmed sequences have not yet been officially named by the IUIS TR-IG Review Committee, but the sequences are publicly available at the OGRDB website [https://ogrdb.airr-community.org/sequences/Human] [42].

Finally, reports were compiled from the VDJbase website of genes that were inferred from AIRR-seq data in the study of Gidoni and colleagues [43], which was chosen because it is a particularly high-quality study of FACS-sorted naïve (CD19^+^ CD27^-^ IgD^+^ IgA^-^ IgG^-^) B cells. The VDJbase pipeline uses the TIgGER genotype inference utility [39], and was used to produce the Gidoni-VDJbase dataset of 99 individual heavy and light chain IG genotypes. These genotypes were additionally reviewed and confirmed against VDJbase haplotyping data, whenever possible.

For a candidate IGHV sequence to be included in the IGH Source Set, at least two independent reports of that sequence were required. Such reports could, for example, be provided by two key studies, or a single key study could be supported by Gidoni-VDJbase AIRR-seq data or Gibson-VDJbase genomic sequence data. A report of a genomic sequence from a particular laboratory was not considered suitable confirmation of a separate report of that sequence by the same laboratory, unless the genomic data was clearly reported from multiple individuals. If the multiple origins of genomic germline sequences were unequivocal, as was often the case with sequences from the Wang, Gibson and Rodriguez studies [26; 28; 46], this was accepted as sufficient confirmation of the reported sequence. Multiple reports from different individuals in the Gidoni-VDJBase AIRR-seq dataset were also considered to be adequate confirmation of the reality of a particular sequence, if the sequence was also supported by a suitable GenBank entry that reported an unrearranged sequence.

For most sequences, key studies provided details of genomic mapping. The general location of other sequences within the genome was inferred by careful analysis of selected AIRR-seq datasets. Expression data is unable to distinguish between identical sequences, such as IGHV1-69*01 and IGHV1-69D*01. In the evaluation of such cases, a sequence was confirmed at a particular genomic location if it had been mapped at that locus by at least one of the key genomic studies and if the sequence had been observed in AIRR-seq data.

### Compilation and evaluation of data relating to IGHD, IGHJ, IGKJ and IGLJ genes

The set of candidate IGHD and IGHJ sequences was compiled by identifying all sequences reported in the historically important genomic studies of Mattila [58], Corbett [59] and Watson [25], as well as from the recent study of Rodriguez and colleagues [46]. IGHD sequences from the Matsuda [44] study were also included. Candidate IGKJ and IGLJ sequences were similarly compiled from studies of the Leder group [60], the Watson group [24] and the Shimizu group [53], as well as from Gidoni-VDJbase AIRR-seq and Gibson-VDJbase genomic sequence datasets.

For a candidate IGHD, IGHJ, IGKJ or IGLJ gene to be included in the Source Sets, it had to be observed in at least two of these key genomic studies, or in a key study and at least one inferred or directly sequenced genotype from the Gidoni-VDJbase or Gibson-VDJbase datasets. Inferred genotypes were manually reviewed to confirm the validity of any candidate sequence that was observed in just a few individuals, or that was present in an individual’s dataset at low frequency.

Gidoni-VDJbase IGHD genotypes are mostly based upon the identification of partial IGHD genes within VDJ rearrangements. Exonuclease trimming of IGHD gene ends means that rearrangements rarely include full-length IGHD genes. This makes it almost impossible to identify some full-length IGHD genes from VDJ rearrangements with certainty. In this study, genotypes based upon AIRR-seq data were used to confirm critical centrally-located gene-defining and allele-defining nucleotides in selected IGHD sequences. Expression data was not used to confirm the four highly similar genes of the IGHD1 gene family (IGHD1-1*01, IGHD1-7*01, IGHD1-14*01 and IGHD1-20*01), the alleles of IGHD2-2 that are distinguished from one another by terminal nucleotides, or the extremely short IGHD7-27 gene. And while expression data can identify the identical IGHD4-4*01/IGHD4-11*01 and IGHD5-5*01/IGHD5-18*01gene pairs, it cannot be used as evidence in support of the existence of any of those individual genes.

### Development of the AIRR-C Reference Sets from the Source Sets

Truncated sequences in the Source Sets have been extended in the AIRR-C IGH_VDJ, IGKappa_VJ and IGLambda_VJ Reference Sets. IGHV, IGKV and IGLV alleles in the Source Sets were first identified that were shorter than other allelic variants of their genes, or shorter than other highly similar genes. This included most IARC-affirmed alleles. The IARC does not usually affirm the terminal nucleotides of a sequence, but their reports may include recommendations regarding the terminal nucleotides, based upon analysis of the gene ends in the AIRR-seq data. Where such recommendations have been made, they were used as suitable extensions. Recent genomic sequences were also used as a source of many extensions [46; 57], and for some sequences, Gidoni-VDJbase AIRR-seq data [43] was reviewed to identify the most likely gene ends. If no evidence of these kinds were available, extensions of one or two nucleotides were based upon the endings of similar sequences.

The AIRR-C Reference Sets also include modifications to deal with exact paralogs. Where a nucleotide sequence was present as two or more identical entries in a Source Set, only a single entry was retained in the matching AIRR-C Reference Set. The names of any exact paralogs are noted in the metadata associated with the retained sequences.

No investigations of functionality were conducted here. The AIRR-C Reference Sets are restricted to those genes that are known to include alleles that are ORFs. Some genes include alleles that are ORFs and other alleles that are not. A small number of sequences in the Reference Sets that are not ORFs are allelic variants of these genes. The Reference Sets may also include some pseudogenes that are non-functional for reasons that lie outside their coding regions. It is noted within the OGRDB database whether or not each sequence is an ORF, and expression data is available via VDJbase.

GenBank entries were found for all IG genes identified in the key studies. GenBank Accession numbers were identified by BLAST searches for the small number of previously-reported sequences that were only identified by reference to VDJbase data. For all but one of these sequences, BLAST searches identified one or more suitable GenBank Accession numbers. The only reported sequence in GenBank matching IGLV2-8*03 was the truncated sequence Y12418. This sequence is therefore reported in the Reference Set metadata. For all other sequences, accession numbers of full-length unrearranged sequences are reported in the sequence metadata, and a single sequence, selected where possible to include full sequences of the flanking regions, was used as the basis of OGRDB annotation. This ‘representative’ sequence is reported in the associated notes.

## Results

We present optimised germline reference sets for IG heavy (IGH_VDJ), kappa (IGKappa_VJ) and lambda (IGLambda_VJ) genes in which evidence in support of each sequence included in the database has been evaluated and strict criteria have been met. The possible consequences of inclusion of truncated sequences in a Reference Set is first demonstrated.

### Demonstration of the consequences of sequence truncations on AIRR-seq analysis

IgBLAST-based annotation of AIRR-seq data sets of six subjects shown by inference to carry either one of the two alleles of IGHV4-38-2 in their genotypes was assessed. When data sets were annotated using IgDiscover and a Reference Set with a version of IGHV4-38-2*01 that had a two nucleotide 3’-truncation, four individuals who only carried the IGHV4-38-2*01 allele were suggested to be heterozygous at the IGHV4-38-2 locus (Figure 2a). About a third of reads were software-annotated as being derived from IGHV4-38-2*02, with all these alignments including at least 1 difference from the germline sequence (data not shown). Extension of the IGHV4-38-2*01 sequence to cover the likely full-length ending of the gene allowed IgDiscover to annotate essentially all reads in these four individuals as being derived from IGHV4-38-2*01 (Figure 2b). In contrast, annotation of reads derived from IGHV4-38-2 in two subjects who through inference had been shown to express only IGHV4-38-2*02 resulted, as expected, in the vast majority of reads being identified as IGHV4-38-2*02. This was independent of the length of the 3’-end of the IGHV4-38-2*01 sequence. There was a slight increase in misalignments to IGHV4-38-2*01 in these datasets, after extension of the IGHV4-38-2*01 sequence. Review of IgBLAST alignments showed these misidentified VDJ rearrangements to have identical sequence identity to both of the alleles, but the reads were assigned to IGHV4-38-2*01 by downstream analysis programs. In summary, truncated sequences in the Reference Set can negatively affect the quality of sequence annotation.

**Figure 2:**
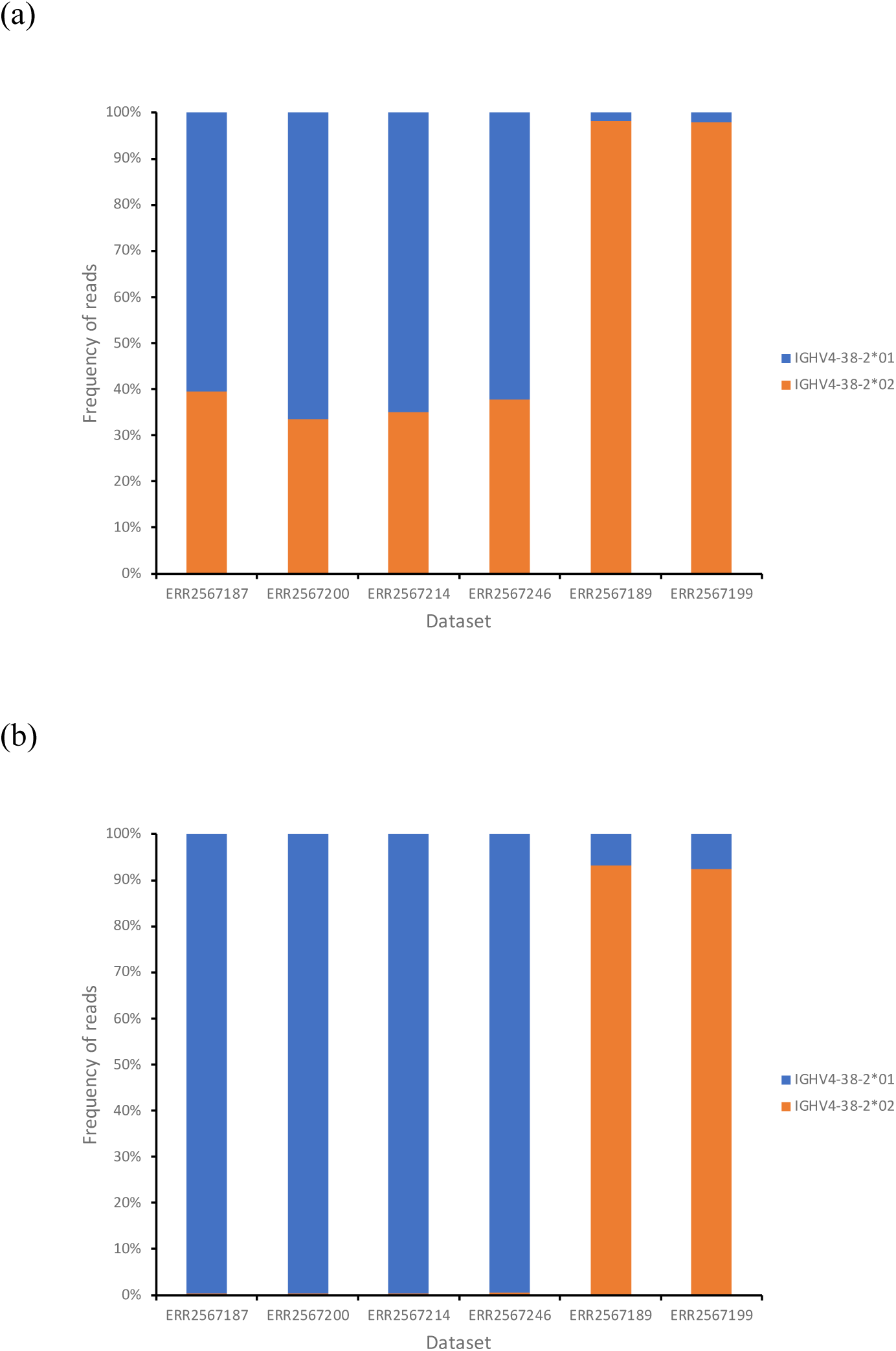
Frequency of alignments to each of two IGHV4-38-2 alleles, before (a) and after (b) the extension of the IGHV4-38-2*01 allele by two nucleotides at its 3’ end. In figure (b), review of the small number of alignments to IGHV4-38-2*01 in the ERR2567189 and ERR2567199 datasets showed them to involve VDJ rearrangements that had identical sequence identity with both of the alleles. In such circumstances, many downstream analytical tools assign the rearrangements to the allele with the lowest allele number.

### Genes of the IGH locus

A set of 295 candidate IGHV gene sequences was compiled from published sources, as outlined. 67 sequences were rejected because they were associated with genes for which no alleles which are ORFs have been seen, because of major truncations, or because the sequences lacked the conserved cysteines. Based on the criteria outlined in the Methods section, we found evidence providing confidence in the existence of 206 of the remaining IGHV sequences, including a number of exact paralogs. The 22 candidate sequences that were not confirmed for inclusion in the IGH Source Set are documented in Supplementary Table I. This will aid their evaluation for possible inclusion in future versions of the Reference Sets, if additional supportive evidence becomes available.

The genomic locations of 167 of the 206 IGHV sequences in the IGH Source Set were confirmed by the key studies or by analysis of Gidoni-VDJbase AIRR-seq data. This analysis focused upon individuals who carried a sequence in question, but also carried other mapped alleles of that particular gene, and were heterozygous at the IGHJ6 locus. Haplotype analysis of AIRR-seq data can show that an unmapped sequence is always associated with a particular J allele, in individuals who are heterozygous at that J locus, while a mapped allele of the same gene is always associated with the alternative J allele (Figure 3). In such cases, the <u>general</u> location of the unmapped sequence can be inferred with confidence.

**Figure 3:**
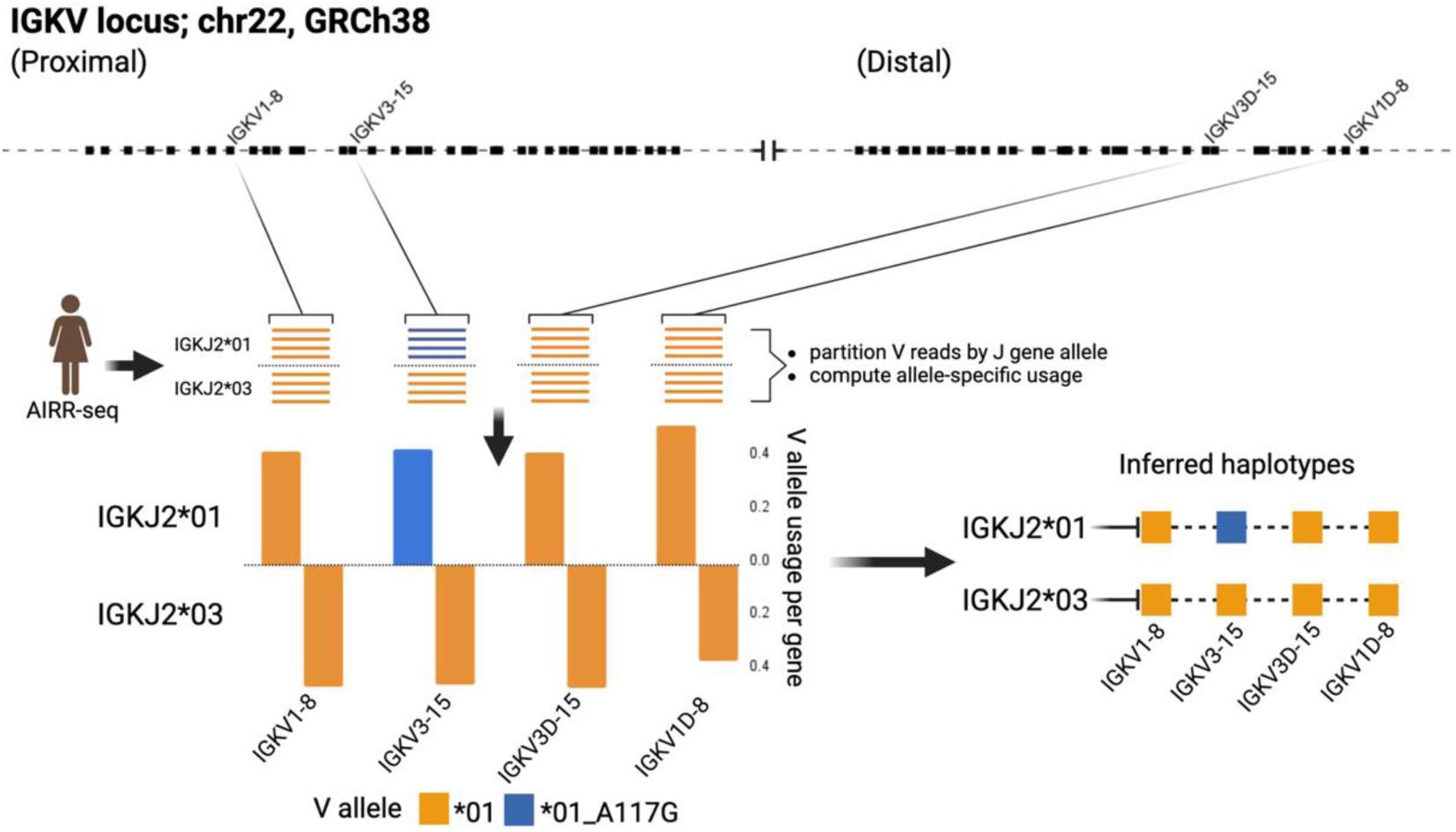
The use of AIRR-seq data to infer the existence and likely location of novel alleles. Data from Gidoni-VDJbase IGK dataset P1_I24 provided strong support for the existence of the novel IGKV sequence IGKV3-15*01_a117g, and of its location in the proximal locus. Inferred haplotypes were determined using the IGKJ2 locus in this heterozygous individual.

Careful review of genomic data [46], numerous AIRR-seq datasets (e.g. Gidoni-VDJBase P1_I10_S1) and previous studies [57; 61] suggest that the IGHV4-59*08 sequence is usually found at the IGHV4-61 locus. As the sequence was observed by Rodriguez and colleagues at the IGHV4-59 locus in one individual [46], the IGHV4-59*08 sequence was included in the IGH Source Set. The identical sequence at the IGHV4-61 locus was given the temporary name IGHV4-NL1*01, and was also included in the Source Set. As the IGHV4-61 locus is the more likely source of the sequence, IGHV4-NL1*01 was selected for inclusion in the Reference Set.

Twenty-nine of the IGHV gene sequences had apparent short 3’ truncations of one or two nucleotides. All these sequences are extended in the IGH_VDJ Reference Set, and this is noted in the metadata. Only one sequence was extended (by two nucleotides) based solely upon the endings of other similar full-length alleles: IGHV5-51*06.

Many pairs of apparently identical IGHV sequences were identified in the Rodriguez study [46]. Twelve alleles of the IGHV1-69 gene, four alleles of the IGHV2-70 gene and three alleles of the IGHV3-23 gene were also observed as exact paralogs at the IGHV1-69D, IGHV2-70D and IGHV3-23D loci [46]. For the moment, this is not reflected in the IGH_VDJ Reference Set, and these observations are not included in counts of sequences in the Source Sets or Reference Sets. Most of these sequences are only included in the IGH Source Set and the IGH Reference Set as alleles of the IGHV1-69, IGHV2-70 and IGHV3-23 genes. Four sequences (IGHV1-69D*01, IGHV2-70D*04, IGHV2-70D*14 and IGHV3-23D*01) are included in the Source Set and are noted as exact paralogs in the metadata associated with IGHV1-69*01, IGHV2-70*04, IGHV2-70*14 and IGHV3-23*01. Evaluation of the genomic location of the other reported IGHV1-69D, IGHV2-70D and IGHV3-23D sequences awaits the publication of assemblies produced using longer genomic sequences.

Four other sequence pairs with names that do not hint at their shared sequence identity are recognized: IGHV3-30*02 and IGHV3-30-5*02; IGHV3-30*04 and IGHV3-30-3*03; IGHV3-30*18 and IGHV3-30-5*01; IGHV4-30-4*09 and IGHV4-31*03. In the IGH_VDJ Reference Set, the IGHV3-30, IGHV3-33 and IGHV4-31 alleles are included, and the exact paralogs at alternative loci are noted in the Reference Set metadata. The data of Rodriguez and colleagues suggests that there are many other paralogs of IGHV3-30, IGHV3-33, IGHV3-30-3 and IGHV3-30-5 genes [46]. These Rodriguez sequences also await formal validation.

Thirty-four candidate IGHD sequences were considered for inclusion in the Reference Set. Three reported IGHD sequences, IGHD3-3*02, IGHD3-10*02 and IGHD3-16*01, were not included as candidate sequences, as they were not reported by any of the key genomic studies and were not identified in the Gidoni-VDJbase IGHD genotypes. Confirmation of most of the candidate IGHD sequences came from their reporting in two or more of the key genomic studies. IGHD2-2*03 was only documented in a single key genomic study and could not be validated with AIRR-seq data. It is therefore not included in the Reference Set. IGHD2-8*02 and IGHD2-21*01 were seen in the genomic sequences of Rodriguez and were confirmed by AIRR-seq alignments. Three candidate IGHD sequences were only seen in the Rodriguez study - IGHD3-10*03, IGHD3-16*03 and IGHD5-18*02. These sequences were unofficially assigned these names when they were recently incorporated into the IMGT database, and they have been included in the Reference Set using these names. Two IGHD gene pairs of identical sequences were seen: IGHD4-4*01 / IGHD4-11*01 and IGHD5-5*01 / IGHD5-18*01. IGHD4-4*01 and IGHD5-5*01 represent the two pairs in the IGH_VDJ Reference Set.

Only eight IGHJ gene sequences were candidates for inclusion in the IGH_VDJ Reference Set, as five other previously reported sequences (IGHJ3*01, IGHJ4*01, IGHJ4*03, IGHJ5*01, and IGHJ6*01) lacked evidentiary support. The reported IGHJ pseudogenes IGHJ1P, IGHJ2P and IGHJ3P were also not considered here. Seven of the eight candidate sequences were confirmed from multiple genomic reports. IGHJ6*04 was only seen in one Gidoni-VDJbase dataset (P1_I69), but as this observation lacked confirmation, the sequence was not included in the IGH_VDJ Reference Set. Recent sequencing data [46; 57] was used to provide the 3’ terminal nucleotide for the IGHJ6*03 sequence in the Reference Set.

### Genes of the Kappa and Lambda Light Chain Loci

102 candidate IGKV gene sequences were identified. Twenty-four sequences were rejected because they were associated with genes for which no alleles which are ORFs that include the conserved cysteines have been seen. Evidence was seen providing confidence in the existence of 71 of the remaining sequences. Seven candidate sequences that were excluded from the Reference Set because of a lack of confirmatory evidence are documented in Supplementary Table II. The genomic locations of all but 8 of the sequences in the Reference Set were confirmed either by direct mapping studies, or by inference from AIRR-seq data. The IGKV inferred mapping analysis focused on individuals who are heterozygous at the IGKJ2 locus. The genomic location of two novel alleles as well as six genes of the duplicated regions of the IGK locus were confirmed with confidence after analysis of Gidoni-VDJbase haplotype-based data (Figure 3). Expression frequency data was also used. Genes of the proximal locus are known to be expressed at significantly higher frequencies than their corresponding genes of the distal locus [50]. The two novel IGKV sequences were confirmed as alleles of IGKV3-15 rather than IGKV3D-15, for like IGKV3-15*01, these sequences were recorded at frequencies greater than 4% of the total repertoire (Figure 4). IGKV3D-15*01 was consistently seen at frequencies of 0.6% or less. The novel sequences were recently affirmed by the IARC and given the names IGKV3-15*i01 and IGKV3-15*i02.

**Figure 4:**
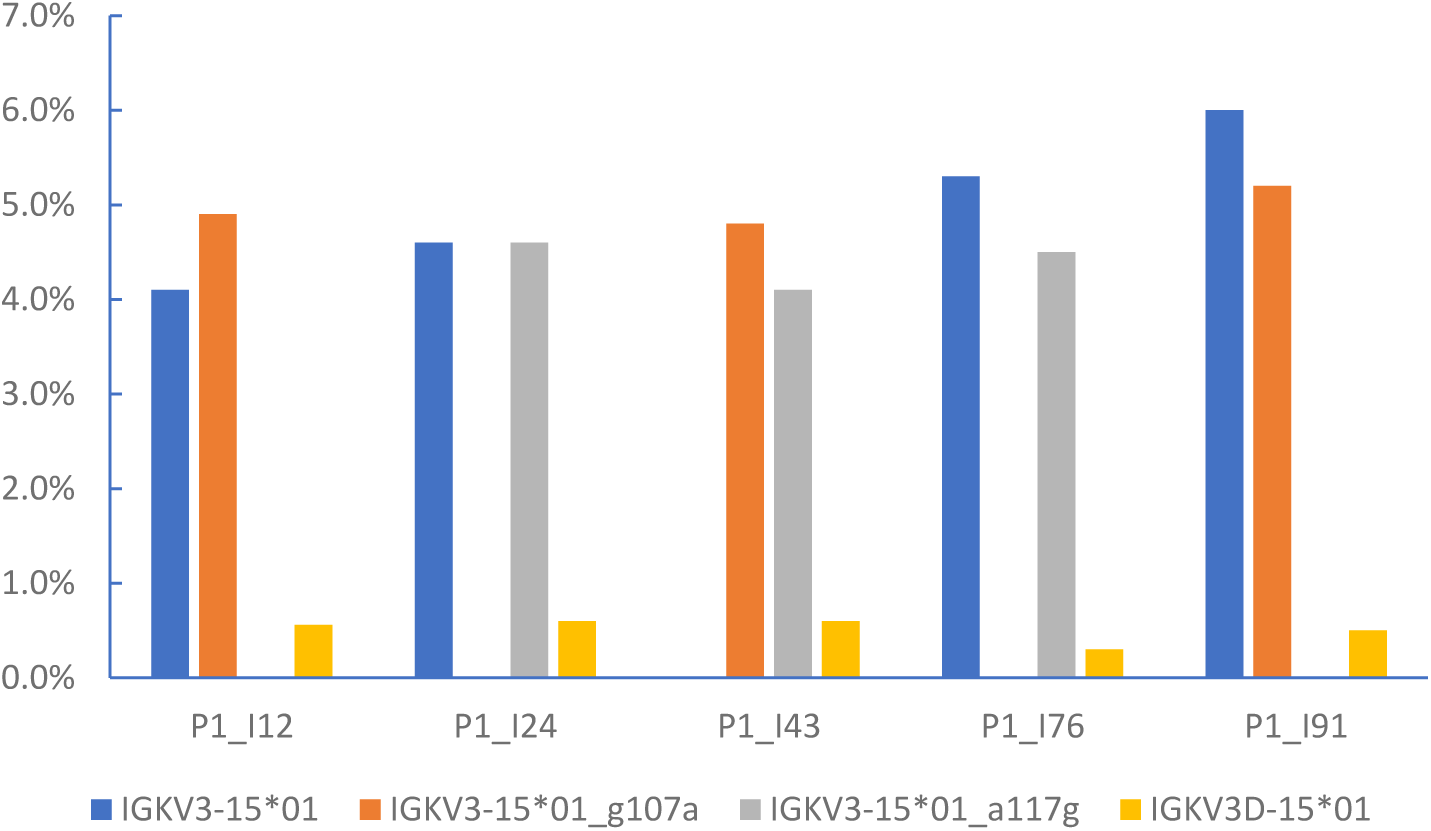
Frequencies of expression of IGKV genes, as percentages of the overall VJ repertoire, in five individuals identified in the Gidoni-VDJbase datasets. These datasets show novel alleles are expressed in three individuals who also express two IGKV3D-15*01 alleles. This suggest that the unmapped novel alleles are located in the proximal kappa locus, and this is also supported by haplotyping of P1_I12 and P1_I24 (data not shown). Expression frequencies give further support for the genomic location of the novel alleles, for genes of the proximal locus are known to be expressed at significantly higher frequencies than genes of the distal locus.

Mapping studies have confirmed that five IGKV sequences of the proximal locus (IGKVare present as exact paralogs in the distal locus. In the IGKappa_VJ Reference Set, these sequences are listed as genes of the proximal locus, with their paralogs noted in the Reference Set metadata.

There were seven candidate IGKJ sequences, and they were all confirmed as suitable for the Source Set, and subsequently for the AIRR-C IGKappa_VJ Reference Set. Two previously reported IGKJ sequences (IGKJ2*02 and IGKJ4*02) were excluded. 71 IGKV and 7 IGKJ sequences therefore form the IGKappa_VJ Source Set, while 66 unique IGKV and 7 IGKJ sequences make up the AIRR-C IGKappa_VJ Reference Set.

115 candidate IGLV gene sequences were first identified, including 5 novel alleles. The novel alleles were all ORFs from the study of Gibson and colleagues [26] that were seen in six or more individuals. Other less-frequently observed novel alleles from the Gibson study await future assessment. Twenty candidate sequences were then rejected because they were associated with genes for which no alleles which are ORFs with the conserved cysteines have been seen. Confirmatory evidence was seen for 81 of the sequences, and the genomic locations of 52 of those sequences were confirmed by the studies used here in the compilation of the set. Haplotype-basedFour mapping was not possible for IGLV genes as there is no commonly heterozygous IGLJ locus by which haplotyping can be performed. Three of the sequences (IGLV2-8*03, IGLV2-14*04 and IGLV3-9*01) had substantial truncations, and in the AIRR-C IGLambda_VJ Reference Set these sequences have been extended by reference to either Gibson-VDJbase or Gidoni-VDJbase datasets. The 14 candidate IGLV sequences that were excluded from the Reference Set because of a lack of confirmatory evidence are documented in Supplementary Table III.

There were ten candidate IGLJ gene sequences and they were all confirmed as suitable for the Source Set. IGLJ2*01 and IGLJ3*01 are identical sequences and in the Reference Set only IGLJ2*01 is shown. Nine IGLJ and 81 IGLV sequences therefore form the AIRR-C IGLambda_VJ Reference Set.

The Reference Sets were compared to the IMGT Reference Directory that is used by the IMGT/V-QUEST alignment utility, focusing on genes for which ORFs are observed and the conserved cysteines are present (Table I). In addition to gene pairs that are only represented once in the AIRR-C Reference Set, a large number of sequences are present in the IMGT Reference Directory but absent from the AIRR-C Set. However, analysis of the Gidoni-VDJbase dataset shows a very small number of alignments to sequences other than those in the Reference Set (Table I).

**Table I:**
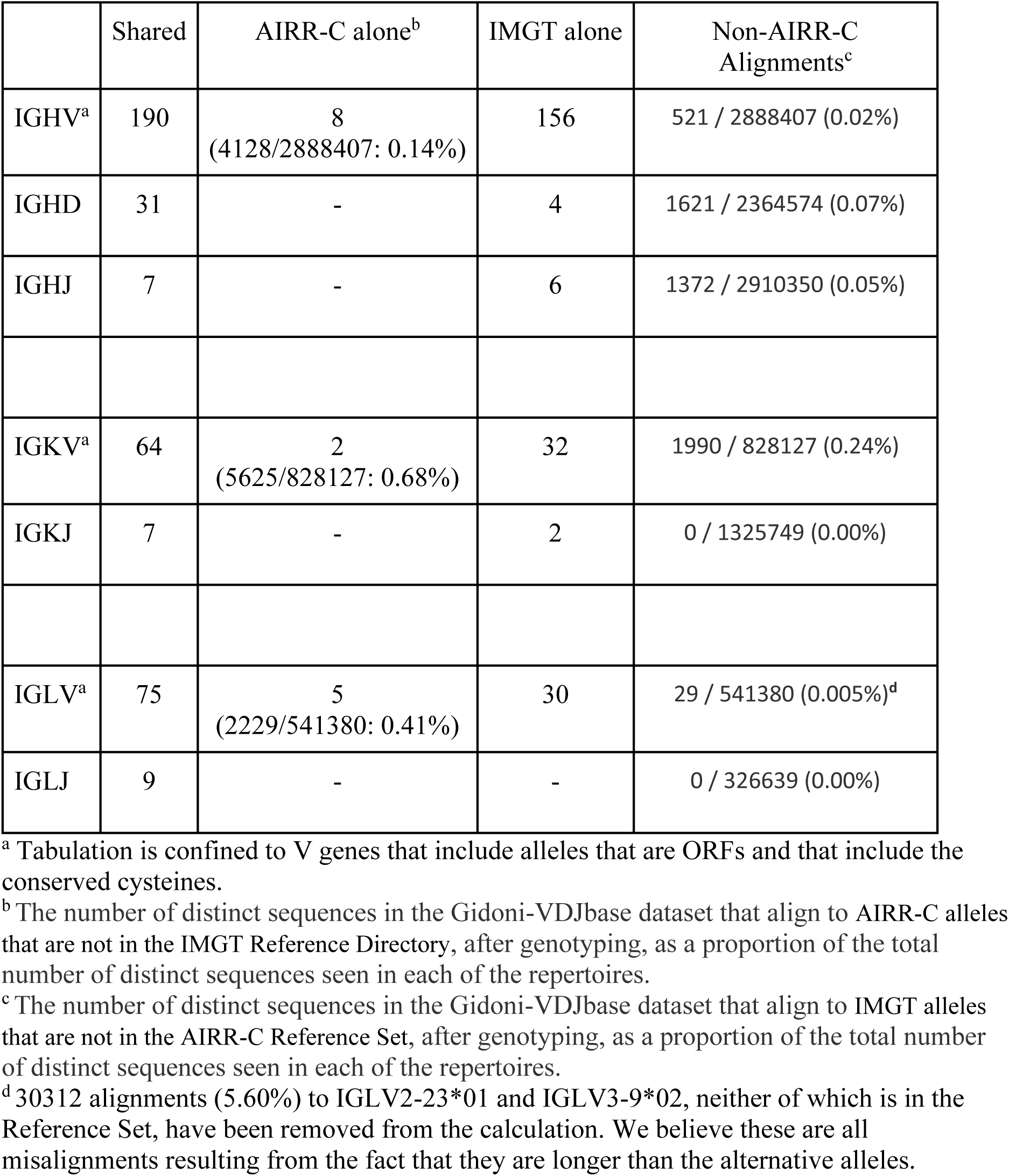
Number of IG alleles that are shared by the AIRR-C Reference Set and the IMGT V-QUEST Reference Directory (accessed 24/07/2023), the number of alleles that are unique to the AIRR-C Reference Set, and the number of alleles unique to the IMGT Reference Directory. Genes that exist as two or more exact copies in the genome are only represented once in the table. Orphons (e.g. IGHV1/OR15*01) are not included in the tabulation.

|  | Shared | AIRR-C alone<sup>b</sup> | IMGT alone | Non-AIRR-C Alignments<sup>c</sup> |
| --- | --- | --- | --- | --- |
| IGHV<sup>a</sup> | 190 | 8<br>(4128/2888407: 0.14%) | 156 | 521 / 2888407 (0.02%) |
| IGHD | 31 | - | 4 | 1621 / 2364574 (0.07%) |
| IGHJ | 7 | - | 6 | 1372 / 2910350 (0.05%) |
| IGKV<sup>a</sup> | 64 | 2<br>(5625/828127: 0.68%) | 32 | 1990 / 828127 (0.24%) |
| IGKJ | 7 | - | 2 | 0 / 1325749 (0.00%) |
| IGLV<sup>a</sup> | 75 | 5<br>(2229/541380: 0.41%) | 30 | 29 / 541380 (0.005%)<sup>d</sup> |
| IGLJ | 9 | - | - | 0 / 326639 (0.00%) |

<sup>a</sup> Tabulation is confined to V genes that include alleles that are ORFs and that include the conserved cysteines.
<sup>b</sup> The number of distinct sequences in the Gidoni-VDJbase dataset that align to AIRR-C alleles that are not in the IMGT Reference Directory, after genotyping, as a proportion of the total number of distinct sequences seen in each of the repertoires.
<sup>c</sup> The number of distinct sequences in the Gidoni-VDJbase dataset that align to IMGT alleles that are not in the AIRR-C Reference Set, after genotyping, as a proportion of the total number of distinct sequences seen in each of the repertoires.
<sup>d</sup> 30312 alignments (5.60%) to IGLV2-23\*01 and IGLV3-9\*02, neither of which is in the Reference Set, have been removed from the calculation. We believe these are all misalignments resulting from the fact that they are longer than the alternative alleles.

## Discussion

Analyses of AIRR-seq data, and even analyses of solitary V(D)J gene rearrangements, generally require the use of germline gene reference sets that are as complete as possible, and that contain as few erroneously reported sequences as possible. These conflicting demands posed a challenge in the preparation of the AIRR-C IG Reference Sets. We have erred here on the side of caution. By restricting our sources of reported germline genes to a small number of key studies, we have likely removed all sequences that were reported in error. Inevitably, some real sequences will also have been lost from the Reference Sets, for the moment, as a consequence.

We know that analysis of new high-quality AIRR-seq expression data and new genomic data should allow most currently excluded but genuine sequences to be confirmed in the near future. Many other novel alleles have already been reported in the studies that were used to establish the Reference Sets [43; 46]. The present study was only able to evaluate a handful of novel sequences for inclusion in the Reference Sets, and the evaluation of other sequences is a task for the future. Even before this happens, it is likely that these first versions of the Reference Sets include nearly all common IGHV, IGKV, IGLV, IGHD, IGHJ, IGKJ and IGLJ allelic variants and many less-common variants that are found in those populations that presently receive the most attention from AIRR-seq researchers.

We believe that a relatively small number of previously-reported sequences that are absent from the Reference Sets are real functional alleles, and many of the ‘missing’ IGHV sequences have previously been flagged as likely resulting from sequencing errors in the 1980s and 1990s when accurate sequencing was particularly challenging [40]. For example, IGHV3-30 is often described as the most polymorphic of the human IGHV genes, but most of the reported IGHV3-30 alleles came from a single study that amplified sequences from a single individual [62]. No additional evidence has ever emerged in support of 14 of these IGHV3-30 alleles, and they are not included in the AIRR-C IGH_VDJ Reference Set. Most of the rejected IGHD and IGHJ genes have also previously been identified as having been reported in error [63; 64].

Other ‘missing’ IGHV sequences have only recently been reported, so they cannot be supported by older key studies. Forty-two human IGHV sequences have been added to the IMGT database in 2023, and 22 of these sequences are now included in the IMGT Reference Directory. The other sequences are described by IMGT as out-of-frame pseudogenes. Only five of the 22 sequences were seen in the Rodriguez study [46], and only four of these sequences are included in the AIRR-C Reference Set. It remains unclear whether the other sequences are rarely carried in the human populations that were the focus of the Rodriguez study, or whether they have been reported in error. In time, if these sequences have been accurately reported, accumulating evidence should lead them to be included in the AIRR-C Reference Set.

Many sequences that IMGT has described as IGHV, IGKV or IGLV pseudogenes are also absent from the AIRR-C IG Reference Sets. The failure to confirm the existence of these pseudogenes may partly reflect the inability of AIRR-seq data to provide such evidentiary support. If they are seen at all in AIRR-seq V(D)J rearrangements, pseudogenes are likely to be very rare. The Gidoni-VDJbase data was therefore generally unable to provide confirming evidence in support of pseudogenes. If real pseudogenes have been omitted for this reason, however, it will have little consequence for AIRR-seq analysis.

The three ‘Not Located’ sequences in the Reference Sets were given additional attention, for some might believe that the failure to map such distinctly different sequences to a genomic assembly is an indication that the sequences were reported in error. IGHV4-NL1*01 was named as part of this report, and should be considered ‘misplaced’ rather than being unlocatable in the genome. We believe this sequence will soon be officially named as an allelic variant of the IGHV4-61 gene.

BLAST searches identified IGKV1-NL1*01 in two FOSMID clones (AC253566 and AC215521) and a BAC clone (AC145029). The novel gene appears to be located between two genes at the far end of the kappa distal locus: IGKV3D-7 and IGKV1-D8. There can be no doubt that this gene exists.

The likely genomic location of IGHV3-NL*01 could not be investigated as it has not been reported from FOSMIDS or BAC clones. In fact, it has only been reported from Papua New Guinean samples. The sequence has been seen in genomic amplifications from buccal swabs, using IGHV RSS-based primers [28], and in cDNA amplified from PBMCs using IG constant region-based primers [65]. The sequence was identified in multiple individuals from samples collected at different places and times [28; 65]. The IMGT database notes that the sequence might be a result of chimeric amplification, though this possibility was ruled out in the original report of the sequence [28]. We strongly believe that the sequence has been accurately reported, and its existence likely points to the significant structural variation that remains to be documented in different human populations.

The AIRR-C Reference Sets are different from other reference sets in several critical ways. Importantly, the reasons why sequences are present in, or sequences have been excluded from the Reference Sets are clearly documented, and the OGRDB website where the Reference Sets can be accessed allows easy interrogation of the data that underpins the reporting of each sequence.

The AIRR-C Reference Sets report whether sequences in the Reference Sets are ORFs, but they do not attempt to identify sequences as being either Functional or Pseudogenes. The functionality of IG genes is not easily defined for there is still no comprehensive understanding of the impact of genetic variation in non-coding regions upon IG expression. There have been some studies of variations in elements such as Recombination Signal Sequences (RSS) [66; 67], but it is unclear, for example, why some apparently functional IGHV genes (e.g. IGHV4-4*01) and IGHD genes (e.g. IGHD6-25*01) are present at such low frequencies in the expressed VDJ repertoire [63; 68] despite these genes reportedly being associated with functional RSS [37]. For this reason, there was no attempt to document functionality in the AIRR-C Reference Sets.

In contrast to other reference sets, the AIRR-C IG Reference Sets have been developed to address specific challenges with AIRR-seq analysis. While the Source Sets report sequences exactly as they have been reported in key studies, or as they have been affirmed from AIRR-seq data by the IARC, the AIRR-C IG Reference Sets include modifications to facilitate accurate analysis of AIRR-seq data.

Many reported sequences are truncated, and this is particularly true for inferred sequences. Although V sequences can be inferred from AIRR-seq data with great confidence for most of their length, the terminal 3’ nucleotides are usually less certain [69]. This is because of data loss resulting from exonucleolytic trimming of gene ends during V(D)J recombination. Other historical reports of germline genes describe sequences that now appear to have short 3’ truncations as a result of problems identifying the boundary between the V gene exons and their RSS. We have demonstrated the consequences of truncations on alignments by analysis of VDJ rearrangements involving the IGHV4-38-2 gene. The consequences are also clear in a review of lambda VJ rearrangements identified as involving IGLV2-23*01 and IGLV3-9*02 when sequences were aligned against the IMGT Reference Directory (Table I). These two sequences are not in the AIRR-C IGLambda_VJ Reference Set, and we believe that all these alignments are in error. We believe that these sequences all involve rearrangements of IGLV2-23*03 and IGLV3-9*01, but these shorter alleles have been overlooked by the alignment utility. In the AIRR-C Reference Sets, apparently truncated sequences have been extended, using IARC recommendations and recent genomic evidence. Extended alleles are listed in the reference set release notes.

The AIRR-C Reference Sets have also been developed to deal with analytical challenges that arise from the presence of exact paralogs in a Reference Set. To allow accurate analysis, the AIRR-C Reference Sets list each duplicate sequence only once. This involves IGHV and IGHD gene pairs, as well as IGKV sequences that are present as identical copies in both the proximal and distal IGKV loci, and a single IGLJ gene pair. The names of all duplicate sequences in the Reference Sets are noted in the reference set metadata and in the reference set release notes. The OGRDB website provides access to a script that facilitates the production of Reference Sets with details of exact paralogs and sequence aliases in the FASTA header (https://airr-community.github.io/receptor-germline-tools/_build/html/introduction.html), should that be desired.

The AIRR-C Reference Sets will provide an avenue by which high quality sequences that lack official names can be rapidly brought to the attention of the research community. This is an issue of growing importance because of an emerging problem with gene reporting. The number of groups of genes that are recognized for their similarities has recently grown, with the documentation of previously unknown structural variation and gene duplication in the immunoglobulin heavy chain (IGH) locus [46]. Until a new approach to the IG nomenclature is developed, there will likely be delays in the official naming of many newly discovered human alleles that appear to belong to these gene groups. The AIRR-C Reference Sets aim to provide early access to such sequences, if their existence is supported by multiple lines of evidence.

Updates to the Reference Sets will be managed by the IARC and the AIRR-C Germline Gene Working Group, and details of future reviews will be publicly available. Only sequences identified in studies of the highest quality will be considered for future inclusion in the sets. If additional inferred sequences are included in the Reference Sets prior to their consideration by the IUIS TR-IG NRC, sequences will be reported using IARC-assigned or other temporary names. These names will be recorded as ‘aliases’ when and if official names are given to the sequences. In the future, it will therefore be possible for reports of each sequence to be easily traced through the literature.

We believe that together, these many features make the AIRR-C Reference Sets far superior to existing publicly available human IG Reference Sets, and the AIRR-C Germline Gene Working Group strongly recommends their use for all human AIRR-seq analysis.

The AIRR-C IGH_VDJ, IGKappa_VJ and IGLambda_VJ Reference Sets, as well as the Source Sets from which they are derived, are available at the OGRDB website. They are published using the AIRR Data Schema [70] which has recently been revised to allow for the definition of Reference Sets [33]. The AIRR-C schema for GermlineSet is supported, and germline sets are downloadable in JSON format compliant with the schema, or in FASTA format. They can also be queried via a REST API. OGRDB manages versioning and change control, such that both users and curators can identify the addition, removal, or modification of sequences in a germline set, and drill down to individual records for each sequence to reveal details. Data published on OGRDB is provided under a minimally restrictive Creative Commons CC0 1.0 licence. OGRDB data is periodically archived at Zenodo (https://zenodo.org) for long-term storage, and each version of a germline set is also deposited at Zenodo and allocated a DOI (https://www.iso.org/standard/81599.html): hence users may cite a persistent identifier that uniquely references the set used in their work.

## Acknowledgements

We thank Christian E. Busse, German Cancer Research Center, Heidelberg, and Ivana Mikocziova, University of Turku, Finland for their valuable comments on earlier drafts of the manuscript.

## Author Contributions

The study was designed by AC, WL, MO, CW, GY, AP, MC and JH. Data analysis was performed by AC, WL, CW, WG, OR, MO, DR and JM-B. The manuscript was prepared by AC, WL, CW, MO and ER. All authors contributed to the refinement of the analyses, the editing of the article and approved the submitted version.

---

**Supplementary Table I:** Evidence in support of the existence of human IGH genes that were candidates for inclusion in the AIRR-C IGH_VJ Reference Set, but which lacked sufficient evidence for inclusion.

| Gene Name | Matsuda<sup>1</sup> | Watson<sup>2</sup> | Watson 2<sup>2</sup> | Scheepers<sup>3</sup> | Wang<sup>4</sup> | Narang<sup>5</sup> |
| --- | --- | --- | --- | --- | --- | --- |
| IGHV1-45*01 | M99645 |  |  |  |  |  |
| IGHV1-69*16 |  |  |  | MG719328 |  |  |
| IGHV2-5*05 |  |  |  |  |  |  |
| IGHV2-26*03 |  |  |  | MG719336 |  |  |
| IGHV2-26*05 |  |  |  |  |  |  |
| IGHV2-70*03 |  |  |  |  |  |  |
| IGHV2-70*13 | AB019437 |  |  |  |  |  |
| IGHV2-70*16 |  |  |  | MG719341 |  |  |
| IGHV2-70*17 |  |  |  | MG719342 |  |  |
| IGHV3-13*02 | M99653 |  |  |  |  |  |
| IGHV3-15*02 | M99654 |  |  |  |  |  |
| IGHV3-16*01 | M99655 |  |  |  |  |  |
| IGHV3-21*04 |  |  |  |  | HM855720 |  |
| IGHV3-25*01 | M99661 |  |  |  |  |  |
| IGHV3-33*02 | M99665 |  |  |  |  |  |
| IGHV3-47*03 | M99674 |  |  |  |  |  |
| IGHV3-49*01 | M99676 |  |  |  |  |  |
| IGHV3-62*01 | AB019437 |  |  |  |  |  |
| IGHV4-28*06 |  |  |  |  | HM855925 |  |
| IGHV4-30-2*03 |  |  |  |  |  |  |
| IGHV4-31*02 | M99683 |  |  |  |  |  |
| IGHV4-34*02 | M99684 |  |  |  |  |  |
| IGHV4-59*13 |  |  |  |  |  |  |
| IGHV4-61*08 | AB019437 |  |  |  |  |  |

<sup>1</sup> GenBank accession numbers of sequences from the studies of Matsuda and colleagues
<sup>2</sup> GenBank accession numbers of sequences from the studies of Watson and colleagues
<sup>3</sup> GenBank accession numbers of sequences from the studies of Scheepers and colleagues
<sup>4</sup> GenBank accession numbers of sequences from the studies of Wang and colleagues
<sup>5</sup> GenBank accession numbers of sequences from the studies of Narang and colleagues
<sup>6</sup> Observed in the study of Rodriguez and colleagues
<sup>7</sup> Dataset from the study of Gidoni and colleagues, available from the VDJbase website
<sup>8</sup> Observed as a single exact match in AIRR-seq data

**Supplementary Table II:** Evidence in support of the existence of human IGKV genes that were candidates for inclusion in the AIRR-C IGKappa_VJ Reference Set, but which lacked sufficient evidence for inclusion.

|  | Zachau<sup>1</sup> | Winter<sup>2</sup> | Watson<sup>3</sup> | Kawasaki<sup>4</sup> | Gidoni<sup>5</sup> |
| --- | --- | --- | --- | --- | --- |
| IGKV1-6*02 |  |  | GRCh38 |  |  |
| IGKV1D-12*02 |  |  | AC243981 |  |  |
| IGKV1D-13*01 |  |  |  | NG_000833 |  |
| IGKV1D-8*03 |  |  |  |  | P1_I74 |
| IGKV2-4*01 | X72814 |  |  |  |  |
| IGKV2D-26*02 |  | Z27499 |  |  |  |
| IGKV3-7*02 | X72812 |  |  |  |  |

<sup>1</sup> GenBank accession numbers of sequences from the studies of Zachau and colleagues
<sup>2</sup> GenBank accession numbers of sequences from the studies of Winter and colleagues
<sup>3</sup> GenBank accession numbers of sequences from the studies of Watson and colleagues
<sup>4</sup> GenBank accession numbers of sequences from the studies of Kawasaki and colleagues
<sup>5</sup> Datasets from the study of Gidoni and colleagues, available from the VDJbase website

**Supplementary Table III:** Evidence in support of the existence of human IGLV genes that were candidates for inclusion in the AIRR-C IGLambda_VJ Reference Set, but which lacked sufficient evidence for inclusion.

|  | VDJbase Genomic<sup>1</sup> | Winter et al<sup>2</sup> | Kawasaki et al.<sup>3</sup> |
| --- | --- | --- | --- |
| IGLV1-40*02 |  | Z22193 |  |
| IGLV1-40*03 |  | Z22192 |  |
| IGLV1-44*04 | P28_I6 |  |  |
| IGLV1-47*03 | P28_I5 |  |  |
| IGLV10-54*06 | P28_I5 |  |  |
| IGLV2-11*02 |  | Z22198 |  |
| IGLV2-8*04 | P28_I3 |  |  |
| IGLV3-1*02 | P28_I3 |  |  |
| IGLV3-22*02 |  |  |  |
| IGLV3-25*01 |  | X97474 |  |
| IGLV4-60*01 |  | Z73667 |  |
| IGLV5-45*05 | P28_I6 |  |  |
| IGLV7-46*05 | P28_I6 |  |  |
| IGLV9-49*02 |  |  | D87016 |

<sup>1</sup> Datasets from the study of Gibson and colleagues, available from the VDJbase website
<sup>2</sup>GenBank accession numbers of sequences from the studies of Winter and colleagues
<sup>3</sup> GenBank accession numbers of sequences from the studies of Kawasaki and colleagues

